# A personalised approach for identifying disease-relevant pathways in heterogeneous diseases

**DOI:** 10.1101/738062

**Authors:** Juhi Somani, Siddharth Ramchandran, Harri Lähdesmäki

**Author notes:** Co-first author; contributed equally to this work (Juhi Somani, Siddharth Ramchandran).

## Abstract

Numerous time-course gene expression datasets have been curated for studying the biological dynamics that drive disease progression; and nearly as many methods have been proposed to analyse them. However, barely any method exists that can appropriately model time-course data and at the same time account for heterogeneity that entails many complex diseases. Most methods manage to fulfil either one of those qualities, but not both. The lack of appropriate methods hinders our capability of understanding the disease process and pursuing preventive or curative treatments. Here, we present a method that models time-course data in a personalised manner, i.e. for each case-control pair individually, using Gaussian processes in order to identify differentially expressed genes (DEGs); and combines the lists of DEGs on a pathway-level using a permutation-based empirical hypothesis testing in order to overcome gene-level variability and inconsistencies prevalent to heterogeneous datasets from complex diseases. Our method can be applied to study the time-course dynamics as well as specific time-windows of heterogeneous diseases. We apply our personalised approach on two longitudinal type 1 diabetes (T1D) datasets to determine perturbations that take place during early prognosis of the disease as well as in time-windows before seroconversion and clinical onset of T1D. By comparing to non-personalised methods, we demonstrate that our approach is biologically motivated and can reveal more insights into progression of heterogeneous diseases. With its robust capabilities of identifying immunologically interesting and disease-relevant pathways, our approach could be useful for predicting certain events in the progression of heterogeneous diseases and even biomarker identification.

**Availability:** The implemented code of our personalised approach will be available online upon publication.

## 1 Introduction

With the increasing affordability of high-throughput technologies, such as microarray and RNA sequencing, genome-wide time-course gene expression data has become one of the most abundant and routinely analysed type of data [Bar-Joseph *et al.*, 2012] for studying and understanding the molecular mechanisms underlying various complex diseases [Menche *et al.*, 2017]. Encapsulating a wealth of information regarding the prolonged or transient expressions of a large set of activated genes [Bar-Joseph *et al.*, 2012], time-course data also helps us understand and model the (multidimensional) dynamics of complex biological systems or phenomena, such as disease progression [Bar-Joseph *et al.*, 2012; Androulakis *et al.*, 2007; Wang *et al.*, 2008]. It offers us the possibility of deciphering the underlying pathophysiologies and systematic evolutions of human diseases [Androulakis *et al.*, 2007]. A prominent goal in such studies has been to identify genes whose expression levels systematically differ between a case (e.g. disease) and a control (e.g. healthy) group, and can be classified as biomarkers for diagnosis and prognosis of the disease.

For more than a decade, various methods have been introduced for modelling time-course data to identify differentially expressed genes (DEGs). Nonetheless, modelling, interpreting and validating the gene expression patterns are continually met with major challenges. The challenges can be largely classified into two categories: (i) robustly modelling the dynamics of time-course data and (ii) accounting for the heterogeneity of complex diseases.

Many methods have been proposed that deal with the most prominent limitations of modelling gene expression time-course data. Some such limitations include non-uniform sampling [Bar-Joseph *et al.*, 2012; Bar-Joseph, 2004], too few sampling times, missing time points, few or no replicates [Bar-Joseph, 2004], autocorrelation between successive time points [Bar-Joseph, 2004; Fischer *et al.*, 2007], and high-dimensionality with small sample sizes [Wang *et al.*, 2008]. Some methods simplify the modelling task by disregarding the dynamic nature and making the expression profiles “coarse-grained” [Wang *et al.*, 2008], such as cross-sectional analysis (i.e. direct time point-wise comparison of samples) [Bar-Joseph *et al.*, 2003] and simplification strategies [Wang *et al.*, 2008; Erdal *et al.*, 2004; Kim and Kim, 2007]. However, these methods are suboptimal. Interpolation methods, such as linear [Aach and Church, 2001] and B-spline (cubic spline) [Bar-Joseph *et al.*, 2003; Storey *et al.*, 2005; Luan and Li, 2004], have been one of the first methods to be attempted for modelling the dynamics of longitudinal data and using them for estimating gene expression levels at unobserved time points [Bar-Joseph *et al.*, 2003; Bar-Joseph, 2004; Fischer *et al.*, 2007]. Even though they incorporate the continuous nature of the data, they may be subject to issues, such as overfitting. In fact, B-spline-based methods require more than ten time points to produce reliable results [Bar-Joseph, 2004; Fischer *et al.*, 2007], which makes it unsuitable for applications in many biological studies [Wang *et al.*, 2008].

Recently, linear mixed models (LMMs) and Gaussian processes (GPs) have become popular choices for time-course data modelling due to their ability of modelling the correlational structure of the data [Verbeke *et al.*, 2010; Wolfinger *et al.*, 2001; Trabzuni *et al.*, 2014]; efficiently handling biological replicates, while accounting for subject-specific variability; including time-invariant and time-varying covariates; and determining the trends over time as well as taking into account the correlation that exists between successive measurements [Kalaitzis and Lawrence, 2011]. Moreover, GP models offer a robust way of estimating missing or unobserved values by providing confidence intervals along the estimated curves of gene expression [Kalaitzis and Lawrence, 2011]. GP models can be used to identify differential expression between multiple conditions [Äijö *et al.*, 2012] or handle general experimental designs [Cheng *et al.*, 2019]. They can also be designed to be robust to outliers and employ flexible model basis [Stegle *et al.*, 2010]. GPs capture the underlying true signal and embedded noise in a time-course gene-expression data in a non-linear manner, without imposing strong modelling assumptions. In addition to answering whether a gene is differentially expressed across the whole time-course, GP models have also been successfully applied for determining specific time-windows when a gene is DE even when no or few observations are made in that time-window [Stegle *et al.*, 2010; Heinonen *et al.*, 2014; Yang *et al.*, 2016].

The traditional applications of these methods detect genes that exhibit different expression levels between a case and a control group (DEGs) across the whole study population. Unfortunately, in the case of heterogeneous data from complex diseases, only a few genes are usually found to be DE across all or most cases because different genes with similar functionalities may be found to be perturbed across cases, thus justifying the gene-level variability at a functional or pathway level [Menche *et al.*, 2017]. In fact, gene-level results from similar studies of heterogeneous diseases, such as cancers [Segal *et al.*, 2004; Drier *et al.*, 2013], asthma, Huntington’s diseases [Menche *et al.*, 2017], rheumatoid arthritis, type 2 diabetes, schizophrenia [Jin *et al.*, 2014], and Parkinson’s disease [Jin *et al.*, 2014; Menche *et al.*, 2017], have often been found to be inconsistent. They show distressingly little overlap between similar studies of the same disease [Subramanian *et al.*, 2005; Segal *et al.*, 2004; Chen *et al.*, 2013; Menche *et al.*, 2017]. Due to these challenges, many methods that summarise the results on a pathway-level have been developed, where the genes are unified under biological themes that aid in a functional understanding of the results. This can be further improved by developing personalised approaches for identifying enriched or disrupted pathways in complex diseases. Here, personalised approach refers to such methods that do not assume that changes are consistent across all study subjects but instead they identify biomarkers for each subject, e.g. by analysing each case-control pair separately; and a pathway is an overarching term for a group of genes unified under biological themes and are also referred to as gene sets in Subramanian *et al.* [2005].

Menche *et al.* [2017] introduced a framework for personalised gene expression analysis, where personalised perturbation profiles (PEEPs) are constructed per case subject by calculating a z-score with reference to the control group and considering any gene with a z-score above an optimised threshold to be part of the PEEP. Using a combinatorial model on the PEEPs, they strive to identify a single pool of disease-associated genes that can be used to accurately predict the disease status of each subject. The method of Menche *et al.* [2017] thus accounts for heterogeneity. However, it is not directly suitable for modelling time-course data.

Pathway (gene set) enrichment analyses, such as Fisher’s exact test and GSEA [Subramanian *et al.*, 2005], are commonly applied to the gene-level results in order to obtain an understanding of the results at the level of biological processes. Several specialised methods have also been proposed for pathway-level analysis with two groups, such as module map [Segal *et al.*, 2004], CORGs [Lee *et al.*, 2008], Pathifier [Drier *et al.*, 2013], SPCA [Chen *et al.*, 2008], and PARADIGM [Vaske *et al.*, 2010]. However, only a few can be applied directly to time-course experiments. One such method is the unified statistical model for analysing time-course experiments at the pathway level using linear mixed effects models [Wang *et al.*, 2009]. This method directly identifies significant pathways expressed over time by using random effects to model the heterogeneous correlations between the genes in the pathway as well as other fixed and random effects. Unfortunately, these methods do not apply a personalised approach for the modelling.

In this paper, we propose a method that models the time-course data in a personalised manner using Gaussian processes and combines the lists of DEGs on a pathway level. Our method assumes an experimental design where each case subject is matched with a carefully chosen control subject, and the method uses a robust yet efficient method to detect DE genes for each individual with respect to the matched control. Individual-specific gene-level results are summarised at pathway-level using a permutation-based empirical hypothesis testing that is tailored for personalised DE analysis. To study expression changes associated with particular time periods, such as time before disease onset, we also extend the method to detect DEGs in specific time-windows.

We applied this method to two type 1 diabetes (T1D) microarray datasets from Kallionpää *et al.* [2014]. There is growing evidence that T1D is a genetically heterogeneous disease [Atkinson *et al.*, 2014; Mukhopadhyay *et al.*, 2018; Tuomi *et al.*, 2014]. Therefore, in order to gain a robust understanding of the molecular mechanisms underlying this complex and heterogeneous disease, one needs to apply a personalised approach on a pathway-level like the one presented here. We report disruptions in pathways during the early progression of T1D (time-course analysis) as well as in the 6 months windows before seroconversion (autoantibody positivity) and clinical diagnosis of T1D. Seroconversion is the time of autoantibody presentation in T1D susceptible individuals and represents the earliest (currently known) signs of disease progression. However, clinical diagnosis of T1D is established at a very late stage of the disease when insulitis has persisted over a long period of time Clark *et al.* [2017]; Pugliese [2017]; ∼ 80-90% of *β*-cells have been destroyed; and hyperglycemia is achieved [Kallionpää *et al.*, 2014; Clark *et al.*, 2017]. Therefore, identifying relevant perturbations at different stages of the disease can help in monitoring and perhaps predicting the significant events in the disease progression. Our personalised approach was able to identify various disease-relevant and interesting pathways from all three analyses, including those that illustrate the intrinsic mechanisms of disease progression. We also compared the results of the proposed personalised approach from the full time-course and time-window analyses with those of a population-wide method as well as the original results from Kallionpää *et al.* [2014]. This method can be applied to other heterogeneous diseases with a similar experimental design and also extended to non-paired case-control datasets.

## 2 Results

### 2.1 Overview of our personalised GP regression and pathway detection method

In this paper, we present a novel and personalised approach for identifying enriched pathways given time-course observations from multiple two-sample (matched case-control) pairs. We apply our method on gene expression microarray data, but the method can be applied to a variety of -omics or other data types. Our method is demonstrated on gene expression time-course datasets with varying number of case/control observations per pair and uneven sampling times (See Section 2.2). We performed three types of analyses using the datasets described in Section 2.2: early disease progression time-course (TC) analysis across the whole study period, time-series analysis within a window before seroconversion (WSC), and time-series analysis within a window before T1D diagnosis (WT1D). We compared the results obtained using our proposed personalised approach in each of the three analyses with those obtained using a combined (non-personalised) method. Figure 1 gives a high-level (b) overview of our analyses and highlights the differences between the two approaches discussed in this paper.

**Figure 1:**
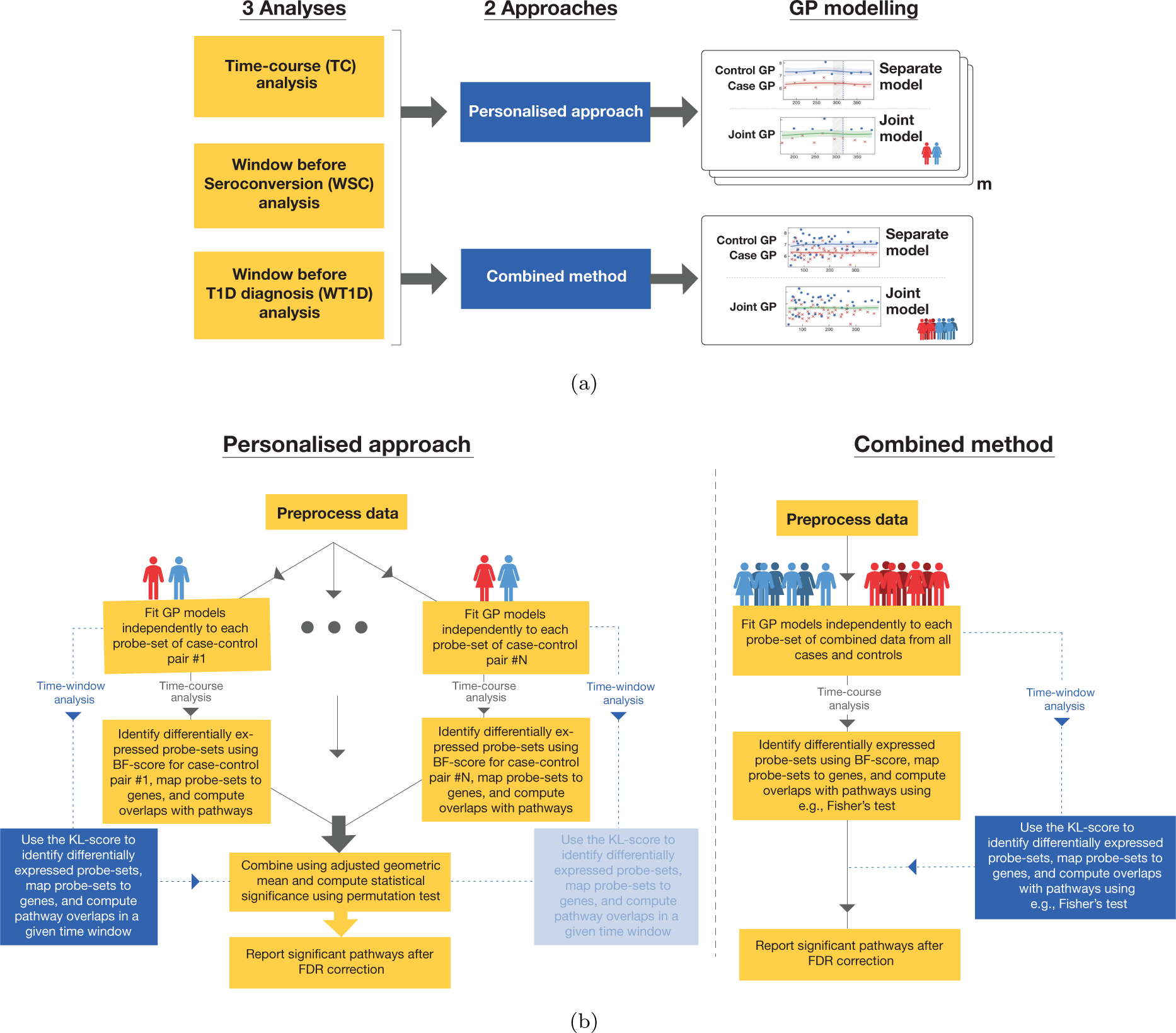
Overview of the method: (a) The three analyses performed using two separate approaches. Here, *m* is the number of case-control pairs. (b) A schematic illustration of our personalised approach and a population-wide approach (combined method). In the personalised approach, we identify DEGs independently for each case-control pair and combine results at the pathway level.

In our personalised approach, we examine each case-control pair and probe-set (i.e. a feature) separately by fitting two models. In the first model, i.e. *joint* model, a GP regression model is fitted to all samples from a case-control pair together (corresponds to the null hypothesis), whereas in the second model, i.e. *separate* model, a case and a control are fitted separately (corresponds to the alternative hypothesis). In the time-course analysis, we identified the differentially expressed (DE) probe-sets for each case-control pair separately by quantifying the fitting for each model using BF-scores (see Equation (6)). For the time-series analyses within a time-window, we derived a method for quantifying differential expression of each probe-set in a specific time-window using KL-scores (see Equation (13)). A probe-set was classified as DE when, in TC analysis, the BF-score is above 4 and, in time-window analyses, the KL-score is above 250. We then map the probe-sets identified as DE from each case-control pair to their corresponding gene names and proceed to perform pathway analysis. Subsequently, we proceed to obtain an enrichment score (over all case-control pairs) for each pathway from the pathway database, MSigDB [Subramanian *et al.*, 2005; Liberzon *et al.*, 2015] by using the metric described in Equation (15). This is followed by a novel permutation test for identifying a set of enriched pathways.

Our personalised approach is significantly different from the combined method where we compute the associated BF-scores and KL-scores per probe-set by pooling together all the cases and all the controls to form a set of combined cases and controls (assuming gene expression difference is homogeneous across the whole study population). The enriched pathways are then identified using a standard one-sided Fisher’s exact test.

We refer the reader to Section 3 for a detailed description of our personalised approach as well as the combined method.

### 2.2 Data

The two microarray datasets used in this study were published by Kallionpää *et al.* [2014]. The raw data (accession code: GSE30211) was downloaded from the GEO database and preprocessed using the *affy*-package in R programming language. Robust multiarray averaging (RMA) normalisation technique was applied on all the downloaded samples.

**Dataset 1** comprised of six case-control pairs chosen from the sample series of *seroconverted progressors*, such that each pair was sampled before and after seroconversion. **Dataset 2** comprised of 15 case-control pairs chosen from the sample series of *T1D progressors*, such that each pair was sampled till at least one month before T1D diagnosis. As subjects in **Dataset 1** were sampled starting before seroconversion, they were used for the early disease progression time-course (TC) and window before seroconversion (WSC) analyses. Whereas, subjects in **Dataset 2** were all sampled after seroconversion and all case subjects progressed to clinical T1D. Hence, they were used for the window before T1D (WT1D) analysis. Here, the case-control pair numbers were kept the same as in Kallionpää *et al.* [2014] for both datasets for comparability.

We use the Molecular Signatures Database (MSigDB, v6.1), which is a collection of annotated gene sets [Subramanian *et al.*, 2005; Liberzon *et al.*, 2015]. We performed pathway-level analyses using 16808 (of 17786) pathways from the collection.

### 2.3 Identifying differentially expressed genes

Differentially expressed genes (DEGs) were identified in a direction-agnostic manner for pathway level evaluation in all three analyses using both the personalised and combined approaches, as explained in Sections 2.1 and 3. In the personalised approach, DEG lists were identified for each case-control pair independently, which resulted in an average of 895, 1127 and 1677 genes DE in the TC, WSC and WT1D analyses, respectively. On average, 14% (TC: 13%, WSC: 13%, WT1D: 17%) of the DEGs overlapped between DEG lists of each case-control pair in the three analyses, thereby demonstrating heterogeneity among case-control pairs. In the combined method, a DEG list was identified by modelling all case-control pairs together for each analysis, which resulted in 436, 234, and 563 genes as DE in the TC, WSC and WT1D analyses, respectively. The overlap of DEGs between the two approaches was significant in all analyses (p-value *<* 0.05 using Fisher’s exact test).

The personalised approach accounts for the heterogeneity between the pairs in time-course and time-window analyses. Firstly, the differential expression of a gene in a case-control pair could be attributed to any of its probe-sets regardless of the probe-set expressed in other pairs. Secondly, the dynamics of gene expression and even the direction of regulation of a DEG is allowed to vary from one case-control pair to another. It is not clear why certain genes behave inconsistently in different individuals, but it could be due to the presence of certain other genes; or any deviation, regardless of the direction, could result in disease-associated perturbation possibly because of the mechanism of regulating the pathway [Menche *et al.*, 2017]. Thirdly, even a gene that is not differentially regulated in most of the case-control pairs can be relevant on the pathway-level. Finally, the GP modelling was able to robustly interpolate over unobserved time points, which was especially important in time-window analyses where sometimes only a few or no samples were available for determining differential expression, as can be seen in Figures 2(a) and 2(b) as well as Supplementary Figures 1 and 2.

**Figure 2:**
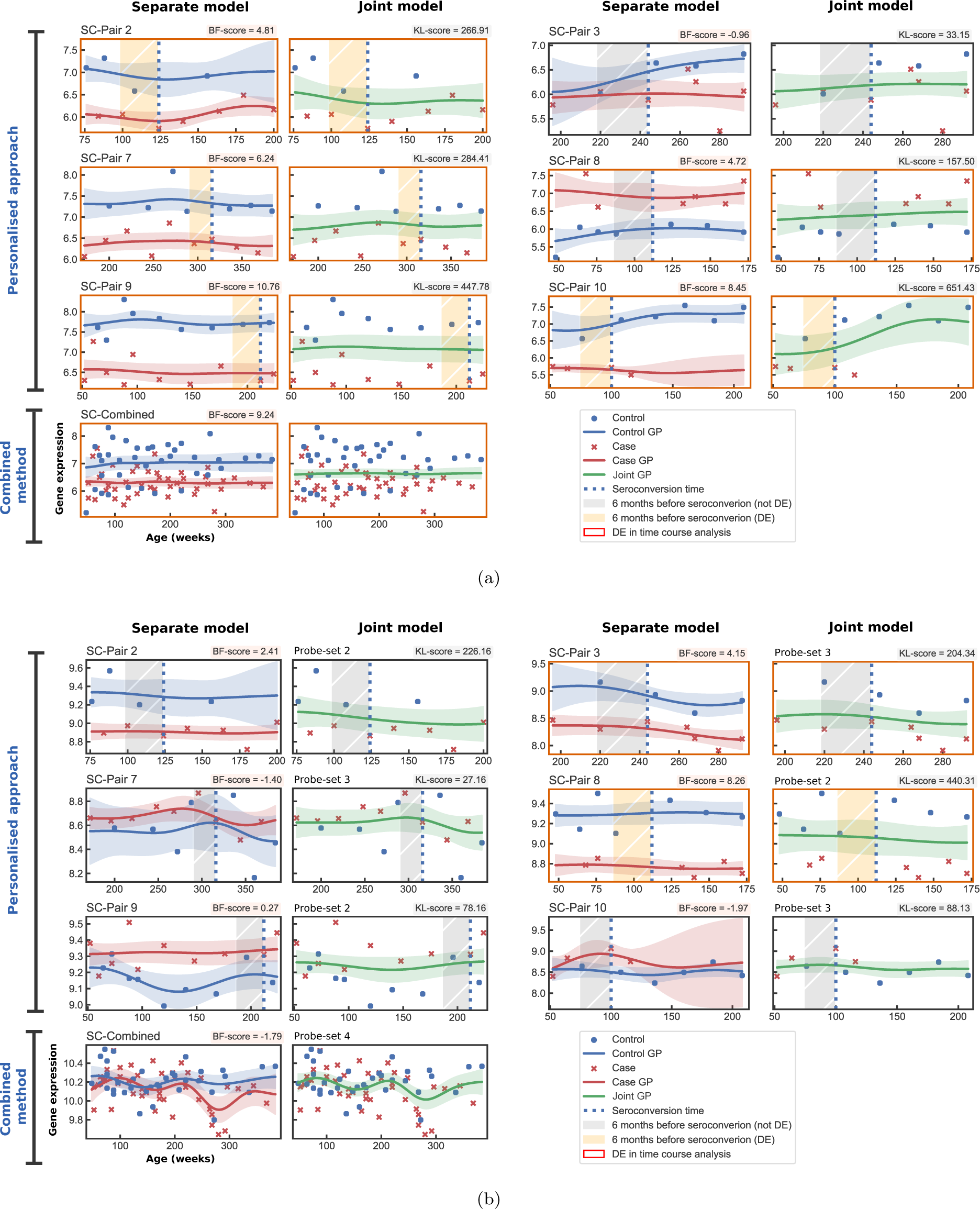
Gene expression plots visualising the GP model fittings of the separate and joint models for the six case-control pairs from **Dataset 1**. A red border around a plot signifies differential expression (DE) in the time-course analysis and an orange shaded window signifies DE in the time-window analysis. Here, pairs from **Dataset 1** are prefixed with ‘SC-’. (a) Gene expression plots for *PTPRN2*. All profiles belong to the same probe-set as all pairs, including the combined method, identified the same probe-set to have the largest BF-score for *PTPRN2*. (b) Gene expression plots for *HSPD1*. The probe-set information is marked for each pair since the profiles identified different probe-sets to have the largest BF-score for *HSPD1*.

The combined method, on the other hand, is more stringent when identifying DEGs in time-course as well as time-window analyses. For a gene to be identified as differentially expressed using this method, one particular probe-set of the gene is usually required to be differentially expressed in almost all of the pairs. Furthermore, if a gene exhibits different temporal expression dynamics or is regulated in opposite directions in different pairs, it is unlikely that this model will identify it as differentially expressed or not.

To illustrate the above-mentioned traits, the expression of the genes encoding the only two autoantigens that were differentially expressed in the TC and WSC analyses, *PTPRN2* and *HSPD1*, from T1D pathway are shown in Figures 2(a) and 2(b). *PTPRN2* encodes a major islet autoantigen in T1D, which plays an important role in insulin secretion in response to glucose stimuli by accumulating normal levels of insulin-containing vesicles and preventing its degradation [Lee, 2019]. *HSPD1* is considered a pro- or anti-apoptotic regulator of apoptosis, depending on the circumstances [Aluksanasuwan *et al.*, 2017], whose high-levels have been associated with diabetes as well as increased expression of inflammatory genes and release of pro-inflammatory cytokines [Blasi *et al.*, 2012; Bellini *et al.*, 2017]. In the TC analysis using the personalised approach, case-control pairs 2, 7, 9 and 10 differentially down-regulated only the *PTPRN2* gene; pair 3 down-regulated only the *HSPD1* gene; and pair 8 down-regulated *HSPD1*, but up-regulated *PTPRN2*. Here, the pairs regulating *HSPD1* differentially express different probe-sets of the gene, whereas all pairs regulating *PTPRN2* differentially express the same probe-set. However, pair 8 up-regulated *PTPRN2* when other pairs down-regulated it. Coincidentally, pair 8 is the only pair that expressed both *PTPRN2* and *HSPD1* in this data and it down-regulated *HSPD1* while up-regulating *PTPRN2*, which may indicate correlation between the two. On the other hand, the combined method found significance only in the *PTPRN2* gene since 5 of 6 case-control pairs differentially expressed the same probe-set. Moreover, Supplementary Figures 1 and 2 show two examples, *HLA_DPB1* (probe-set: 11760799_x_at) and *IRF5* (probe-set: 11726687_a_at), where the case-control pairs regulate the genes in inconsistent directions. Here, the combined method identifies *HLA_DPB1* as DE, whereas *IRF5* is classified as insignificant. The personalised approach, however, identifies both of these genes as significant in all pairs.

### 2.4 Combined method vs personalised approach

Using the combined method, 52, 10 and 80 pathways were found to be significantly enriched with FDR *<* 0.1 in the time-course analysis (TC) of **Dataset 1**, time-series analysis of 6 months time-window before seroconversion using **Dataset 1** (WSC) and time-series analysis of 6 month time-window before diagnosis of clinical T1D using **Dataset 2** (WT1D), respectively. Similarly, 124, 307 and 2550 pathways were found to be significantly enriched with FDR *<* 0.1 in the TC, WSC and WT1D analyses, respectively, using the personalised approach (Figure 3(a), Supplementary Table 1). Of these, 12, 1 and 38 enriched pathways overlapped between the two approaches in the TC, WSC and WT1D analyses, respectively, which was found to be a significant amount (Fisher’s combined p-value *<* 0.0001 obtained from p-values determined using Fisher’s exact test) (Figure 3(b)). Nonetheless, the combined method was unable to identify most of the immunologically interesting and disease-relevant pathways in all three analyses that were identified using the personalised approach.

**Figure 3:**
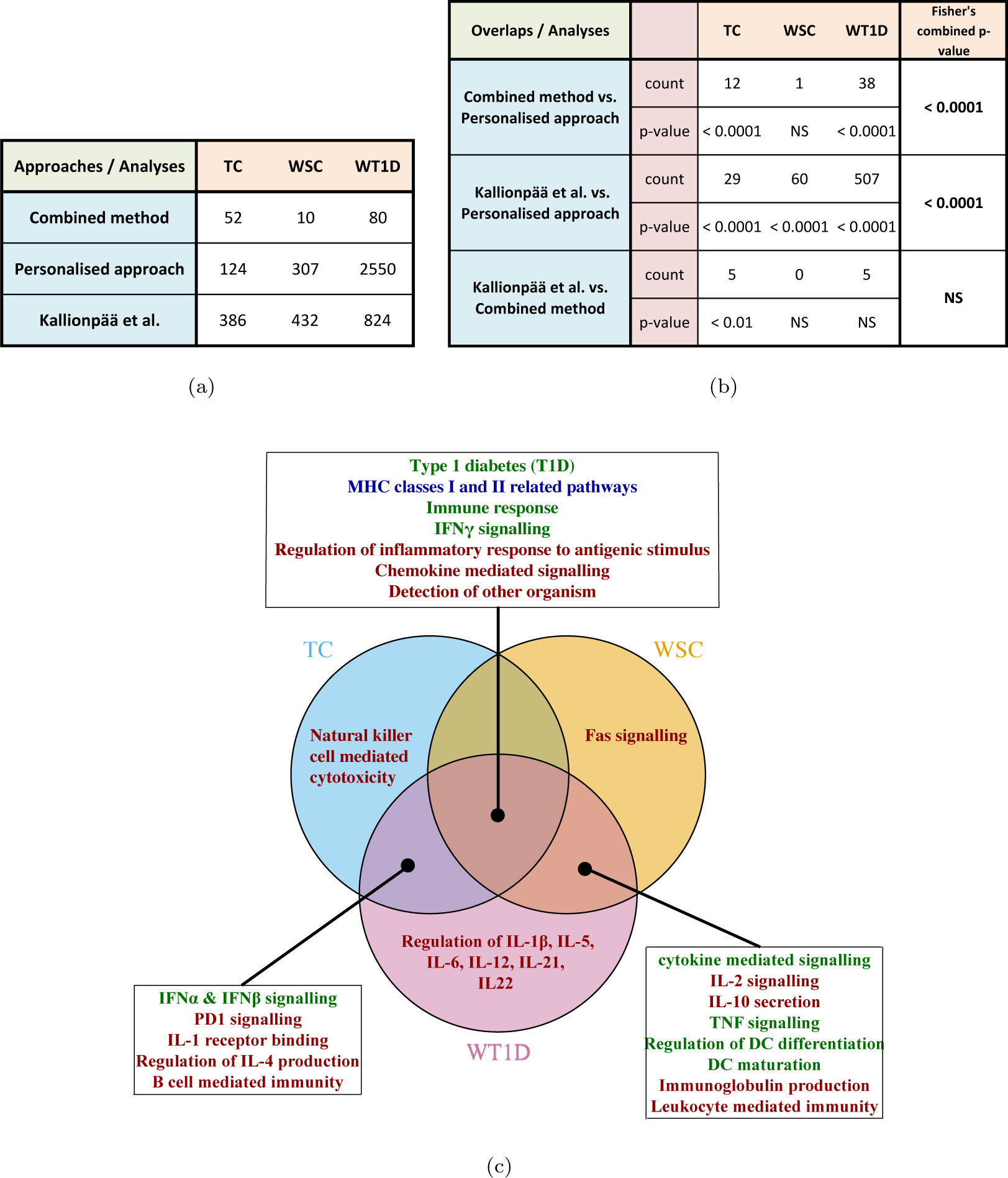
(a) Table listing the number of pathways identified as enriched (FDR *<* 0.1) in TC, WSC and WT1D analyses using the personalised approach, combined method and Kallionpää *et al.* [2014] gene-level results. (b) Table listing the number of enriched pathways overlapping between different approaches in the TC, WSC and WT1D analyses (rows marked as ‘count’). A p-value, determined using Fisher’s exact test, is also given for each overlap to show its significance (rows marked as ‘p-value’), where NS refers to ‘not significant’ p-values (i.e. p-value *>* 0.05). Fisher’s combined p-value over all analyses per comparison is given in the last column. (c) Venn diagram illustrating disease-relevant pathways specific to a certain analysis or overlapping between analyses using the personalised approach. Here, pathways in blue refer to those found enriched by the personalised approach, combined method as well as Kallionpää *et al.* [2014]; green refer to those found enriched by the personalised approach and Kallionpää *et al.* [2014]; and red refer to those found enriched by only the personalised approach; respectively. Full lists of enriched pathways are found in Supplementary table 1.

Among the overlapping pathways, the most disease-relevant pathways were those related to MHC classes I and II protein complexes, protein antigen binding and receptor activity. Where the personalised approach identified the relevance of these pathways in all three analyses, the combined method identified them as significant only in the TC analysis. Moreover, the combined method failed to identify the overarching pathway, ‘antigen processing and presentation’, as significant, which was found to be significant in all three analyses using the personalised approach. Additionally, other interesting and relevant pathways that were identified by the personalised approach were not found using the combined method in any of the analyses.

In particular, the combined method was also unable to identify one of the most basic pathways related to immunological diseases, ‘immune response’, or any of its related pathways in any of the analyses. In fact, the ‘Type 1 diabetes’ pathway was also not found as significant in any of the analyses using the combined method. On the contrary, the personalised approach found the ‘immune response’ pathway as highly significant in all three analyses and many related pathways in at least one analysis. It also identified the T1D pathway as highly enriched in all three analyses.

### 2.5 Enriched pathways identified by the personalised approach

Several disease-relevant and intriguing pathways were identified as enriched using the personalised approach in either all three analyses, only two analyses or uniquely in one analysis. In order to establish relevance of these results, they were cross-validated with the results from the article that published **Datasets 1** and **2**, Kallionpää *et al.* [2014]. The differentially expressed lists of genes from the analyses in the article corresponding to our TC, WSC and WT1D analyses were subjected to a Fisher’s exact test using the pathways from the MSigDB [Subramanian *et al.*, 2005; Liberzon *et al.*, 2015] database to ensure comparability. Their gene-level results identified 386, 432, and 824 pathways as enriched in the TC, WSC, and WT1D analyses, respectively (Figure 3(a)). These pathways overlapped significantly (p-values *<* 0.0001 using Fisher’s exact test) with the enriched pathways found by the personalised approach in all analyses as well as the enriched pathways identified in the TC analysis using the combined method (p-value *<* 0.01), but overlapped insignificantly with the results from time-window analyses using the combined method (Figure 3(b)). Essentially, Kallionpää *et al.* [2014] were able to identify many of the significant pathways identified using the personalised approach. However, Kallionpää *et al.* [2014] mostly identified only the overarching pathways, but not the related pathways with more specialised functions. In some cases, they identified the significance of certain pathways in different analyses than the personalised approach. For instance, the T1D as well as MHC classes I and II related pathways were found enriched (FDR *<* 0.05) in only the WT1D analysis, whereas our method found it in all three analyses. The interesting pathways discussed below that were identified using the personalised approach is illustrated in Figure 3(c) and those identified by Kallionpää *et al.* [2014] and the combined method are highlighted with different colours.

The personalised approach identified significant (FDR *<* 0.05) pathways related to immune response, interferon-*γ* (IFN*γ*) signalling, regulation of inflammatory process to antigenic stimulus, chemokine mediated signalling, and detection of other organism, in all three analyses suggesting their relevance at all stages of the disease (see Supplementary Tables 1). Of these, Kallionpää *et al.* [2014] only identified immune response and IFN*γ* signalling related pathways as enriched (FDR *<* 0.05) in all analyses and detection of other organism pathway was found enriched (FDR *<* 0.05) in only the WT1D analysis.

Multiple interesting overarching pathways were identified as enriched by the personalised approach uniquely in the time-windows right before seroconversion and T1D diagnosis, which were also found by Kallionpää *et al.* [2014] in at least one of the analyses. These include the pathways related to cytokine mediated signalling, TNF signalling, regulation of dendritic cell (DC) differentiation, and DC maturation. However, in contrast to Kallionpää *et al.* [2014] results, the personalised approach was also able to highlight specific cytokine pathways that could be involved in the cytokine mediated signalling as well as possible pathways necessary to regulate/conduct the immune response. In particular, IL-2 and IL-10 related pathways were enriched along with immunoglobulin production, and leukocyte mediated immunity.

Intriguingly, the personalised methods found several pathways that were uniquely enriched during the early prognosis of T1D and in the 6 months window before T1D diagnosis. While IFN*γ* signalling was found significant at all stages of the disease, interferon-*α* (IFN*α*) and interferon-*β* (IFN*β*) signalling were enriched only in the TC and WT1D analyses using the personalised approach, whereas Kallionpää *et al.* [2014] associate their relevance at all stages. Additionally, we found other T1D-associated pathways, such as PD1 signalling, IL-1 receptor binding, regulation of IL-4 production and positive regulation of B cell mediated immunity, to be enriched in the TC and WT1D analyses. However, Kallionpää *et al.* [2014] were unable to detect them.

Furthermore, distinct disease-relevant pathways were determined as uniquely enriched before seroconversion, before T1D diagnosis or during the early stages of T1D progression using only the personalised approach. Specifically, pathways related to natural killer cell-mediated cytotoxicity and Fas signalling were found to be uniquely significant during the early stages of T1D progression and before seroconversion, respectively. Most strikingly, pathways regulating the production of multiple different pro- and anti-inflammatory cytokines, such as Interleukin-1, -1*β*, -2, -4, -5, -6, -10, -12, -21, -22, as well as the related overarching pathways, were found enriched in the 6 months before clinical onset of T1D, where more than half of the cytokine pathways were unique to this time-window.

### 2.6 Type 1 diabetes pathway

The type 1 diabetes pathway was found enriched in all three analyses using the personalised approach. However, the combined method did not find it significant in any of the analyses and Kallionpää *et al.* [2014] found its significance only in the late stages of the disease, i.e. window before clinical onset of T1D. Figure 4 shows the genes that were identified as differentially expressed (BF-score *>* 4) in each analysis per case-control pair (coloured dots). These figures clearly illustrate that only a small fraction of the pathway’s genes are differentially expressed (DE) in most of the case-control pairs and only a subset of these genes are DE in each child. Moreover, the subset of DE genes varies from one pair to another. It is not clearly understood how the presence of certain genes influence that of the other genes, therefore it is not easy to predict which genes in a pathway are selectively or necessarily expressed. When the T1D pathway genes were functionally divided into 3 main sub-processes: release and presentation of autoantigens; activation of CD4+, CD8+ T cells and macrophages; and apoptosis of *β*-cells, it was noticed that at least one gene from each sub-process was identified as DE in each pair. Some pairs did not differentially express any of the (auto)antigen encoding genes, which could indicate an environmental source of (auto)antigens instead of genetic. Similar phenomena may be expected from most other pathways. As an additional example, IFN*γ* signalling pathway has been depicted in Supplementary Figures 3 and 4.

**Figure 4:**
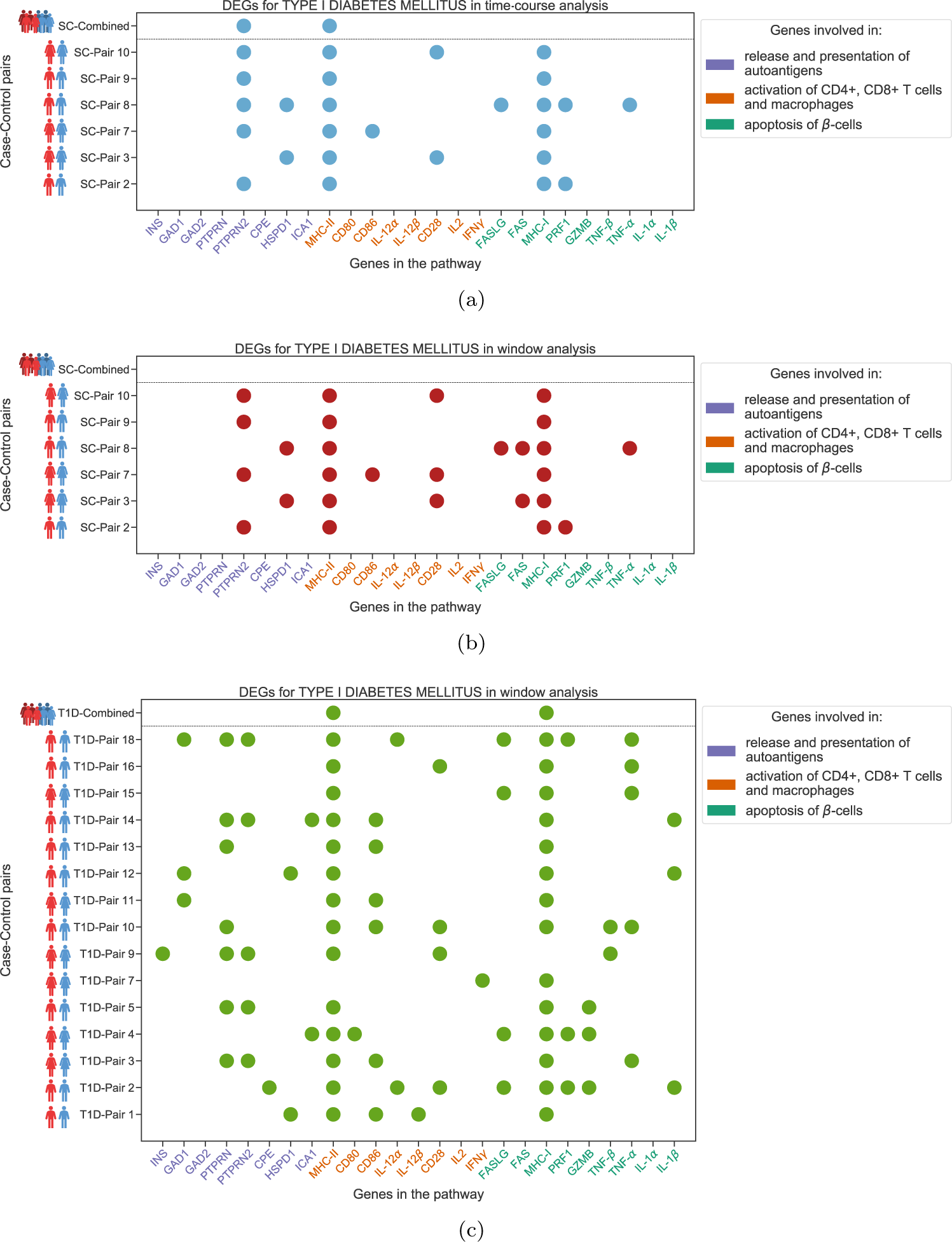
A comparative visualisation of the DEGs between the two approaches for the T1D pathway: (a) TC analysis using **Dataset1**, (b) WSC analysis using **Dataset 1** and (c) WT1D analysis using **Dataset 2**. A coloured dot signifies that the gene is DE in the corresponding case-control pair. Here, the HLA genes from MHC classes I and II are not marked individually, but grouped into their two major classes for convenience; and a class is shown as DE in a case-control pair when at least one probe-set from any HLA gene of the class was found DE in that pair. Also, pairs from **Dataset 1** and **Dataset 2** are prefixed with ‘SC-’ and ‘T1D-’, respectively.

The combined method identified only those genes as DE that were DE in almost all the pairs (Figure 4). Therefore, for a pathway to be recognised as enriched using the combined method, a significant number of the genes in the pathway would need to be DE in most of the pairs, which may not be how heterogeneous diseases, such as T1D, affects pathways.

## 3 Methods

### 3.1 Gaussian process regression

A Gaussian process is a generalisation of the Gaussian distribution. It can be seen as defining a distribution over functions and inference taking place directly in the space of functions [Rasmussen and Williams, 2006]. We denote 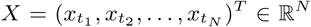 as a vector of noisy measurements for a particular probe-set, which were measured at *N* time points, *T* = (*t*_1_, *t*_2_,…, *t*_*N*_). The GP is defined as

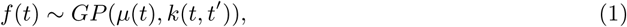

which represents a distribution over function samples *f* (*T*) = (*f* (*t*_1_), *f* (*t*_2_),…, *f* (*t*_*N*_)). Here, *µ*(*t*) is the mean which we assume as zero and *k*(*t, t*′) is a positive semi-definite kernel function, which has kernel parameters *θ*, i.e. *k*(*t, t*′|*θ*). We assume additive Gaussian observation noise *E*, where Gaussian observation is defined as

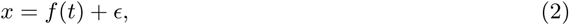

where 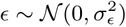 Gaussian process regression modelling involves placing a Gaussian prior, *f* (*T*) ∼*𝒩* (0, *K*_*T,T*_ (*θ*)) over the true model, where the elements of the covariance matrix are defined by the kernel [*K*_*T,T*_ (*θ*)]_*i,j*_ = *k*(*t*_*i*_, *t*_*j*_|*θ*). Here, we use the popular squared exponential kernel, which is defined as

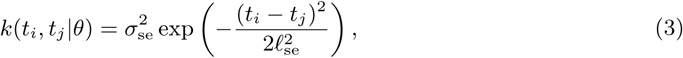

where *ℓ*_se_ is the length-scale parameter that controls the smoothness and 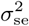 is the signal variance of the kernel. Hence, the kernel parameters are 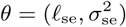.

Given the observed data *X*, the measurement time points *T* and test time points *T*_*\**_, we obtain the posterior distribution *f* (*T*_*\**_) | *X* ∼*𝒩* (*µ*_*\**_, Σ_*\**_) defined by

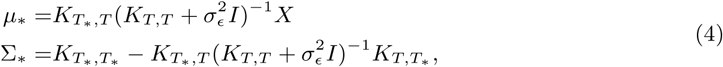

where we denote *K*_*T,T*_ = *K*_*T,T*_ (*θ*) for brevity and 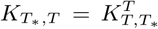 encodes the cross-correlations between measured and test time points.

### 3.2 Prior specification

The gene expression data is first centred to zero by subtracting the mean of the data for GP fitting. This is done independently for the case, control and pooled (case and control) data. For the length-scale (*ℓ*_se_) parameter of the squared exponential kernel, we specify a Gaussian prior (*µ* = 30, *σ*^2^ = 6). We chose the value of *µ* to correspond to 30 weeks which results in a small probability of short length scales and provides a reasonable range of feasible length scales. The magnitude 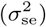 parameter is assigned a square root student-*t* prior (*µ* = 0, *σ*^2^ = 1 and *v* = 20). The noise variance parameter is assigned a scaled inverse chi-square prior (*σ*^2^ = 0.01 and *v* = 1) to restrict it to smaller magnitudes. We use the same (hyper)parameter priors for the case, control as well as joint GPs as explained later.

### 3.3 Marginal likelihood estimation using central composite design

We make use of the central composite design (CCD), which is a form of numerical integration approximation for posterior prediction as proposed in Rue *et al.* [2009]; Vanhatalo *et al.* [2010], to approximate the marginal likelihood (ML). Computing the exact ML is computationally intractable due to the marginalisation over the hyperparameters. Another approach to solving this problem, would be to simply maximise the ML with respect to the hyperparameters. Such an approximation is known as the type II maximum likelihood (ML-II) and can lead to overfitting [Rasmussen and Williams, 2006], especially for small sample sizes common in biomedical studies. Moreover, in our analysis, the ML-II approach failed to generate satisfactory estimates in many instances. CCD assumes a split-Gaussian posterior for log-transformed hyperparameters and defines a set of *R* points 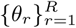 (fractional factorial design, the mode, and so-called star points along whitened axes) that allow for estimating the curvature of the posterior distribution around its mode (see [Rue *et al.*, 2009; Vanhatalo *et al.*, 2010]). We estimate the ML by using the *R* CCD points that are located around the high-probability region of the posterior (which is the integrand in the ML integral) but by replacing the split-Gaussian approximation used for posterior predictions with the exact product of likelihood and prior. In other words, we take the weighted sum of the posterior probability evaluated at the *R* points of the hyperparameter, which are weighted by the integration weights. For a model *M* with data *X*, the estimated ML is given by

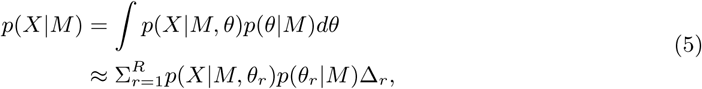

where 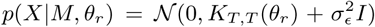 and Δ_*r*_ is *r*^*th*^ integration weight that corresponds to the volume of hyperparameter space allocated to the *r*^*th*^ point. The obtained estimated ML for each model is then used to compute a Bayes factor score, which is used for model selection and for identifying differentially expressed genes (DEGs) as discussed below.

### 3.4 Personalised approach to identifying DEGs in time-course analysis using ML ratio

To identify if a probe-set is differentially expressed between a matched case-control pair, we fit a *joint* and *separate* model to the expression data and identify which model better explains the observed data. The *joint* model involves fitting a Gaussian process over all the data points (i.e. case and control data), whereas the *separate* model involves independently fitting a GP to only the data points corresponding to the cases and fitting another GP to only the data points corresponding to the control. After the fitting, model selection is performed to choose between the *joint* and *separate* model. If the *joint* model is chosen, we conclude that the case and control expressions for the specific probe-set comes from the same process and hence is not differentially expressed. Alternatively, if the *separate* model is chosen, we conclude that the case and control expressions for the corresponding probe-set comes from different processes and hence is differentially expressed. Assume two independent models, *M*^*A*^ and *M*^*B*^, which are fit to the case and control time-course of a particular probe-set, ***x***^***A***^ and ***x***^***B***^, respectively. Also, let a *joint* model, *M*^*S*^, be fit to the pooled data ***x***^***S***^ = (***x***^***A***^; ***x***^***B***^). A standard statistical test would compare models *M*^*A*^ and *M*^*B*^ (*separate* models) against the *joint* model, *M*^*S*^. Hence, the null hypothesis would correspond to no differential expression and the alternate hypothesis would correspond to the presence of differential expression [Stegle *et al.*, 2010].

To perform model selection, we shall compute a Bayes factor score and specify a threshold for a probe-set of a case-control pair to be differentially expressed. The Bayes factor score is calculated as the log ratio of the marginal likelihoods of the *separate* and *joint* models,

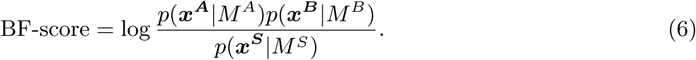

This gives us a score for quantifying the differential expression of each probe-set. We use a threshold of 4 (≈ 54.598 in the linear scale), which corresponds to strong evidence for rejecting the null hypothesis as stated in [Kass and Raftery, 1995]. Probe-sets with BF-score greater than 4 are considered as differentially expressed. The BF-score in Equation (6) is computed for each probe-set and case-control pair separately.

Finally, we map the probe-sets to their corresponding gene names. In case of multiple probe-sets mapping to the same gene name, we choose the probe-set with largest BF-score to represent the gene. This is done independently for each case-control pair, which allows the flexibility of choosing different probe-sets between pairs to represent the same gene.

### 3.5 Personalised approach to identifying DEGs in time-window analyses using KL divergence

In addition to analysing the entire time-course, we also perform a complimentary analysis where we focus on the time-window prior to a significant event (e.g. seroconversion and clinical onset of T1D) to detect disruptions in the pathways. This approach could potentially be used to identify the pathways that are affected before an important event in the prognosis of a disease and hence, can have applications in predictive medicine. In this approach, we propose to detect significant genes by comparing the expression levels of probe-sets between each case-control pair in a 26 week (i.e. approx. 6 months) time-window prior to the seroconversion event and clinical disease onset. The size of the time-window can be chosen as any appropriate duration. We compute the posterior mean and variance of the latent variables of the Gaussian processes within the 26 week time-window, as described in Equation (4), using the representative points of the hyperparameters. We then compute the weighted sum for the mean and variance weighted on the approximative posterior and the integration weights (standard CCD approach for posterior prediction),

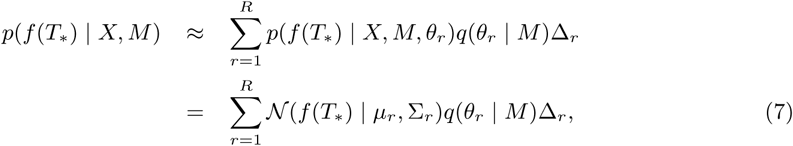

where *µ*_*r*_ and Σ_*r*_ are, as in Equation (4), evaluated with hyperparameter value *θ*_*r*_; *q*(*θ*_*r*_ | *M*) is the split-Gaussian approximative posterior; and *T*_*\**_ defines a time discretisation for the 26 week time interval (we use 26 time points, i.e. a resolution of one week). As prediction with each hyperparameter value *θ*_*r*_ is Gaussian, the combined prediction as a weighted sum is also a Gaussian. Comparisons for the time-window predictions between *separate* (comprising of separate GP fittings for the cases and controls) and *joint* (single GP fitting the pooled case and control data points) models can be made by comparing the distributions using the Kullback–Leibler (KL) divergence [Kullback and Leibler, 1951]. The Kullback-Leibler divergence for any two distributions, *𝒫* and *𝒬*, can be defined as

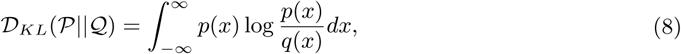

where *p* and *q* are the corresponding densities. To examine the expression level of a probe-set in the time-window, we compare the predictive distributions (Equation (7)) for the *joint* model against the *separate* model in the time-window, by calculating a continuous score obtained using the symmetric KL divergence,

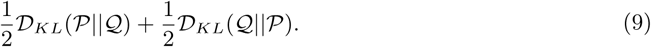

Therefore, to compute the symmetric KL divergence between the *separate* and *joint* model, we assume two multivariate normal distributions: one for the *separate* model, represented by _0_ (previously denoted by *M*^*A*^ and *M*^*B*^); and one for the *joint* model, represented by ℳ_1_ (previously denoted by *M*^*S*^) with dimension equal to twice the number of weeks in the time-window. In the *separate* model, *ℳ*_0_, let 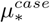 and 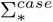 be the predictive mean and covariance matrix (from Equation (7)) for the case GP with the test points taken weekly from the first to the last week of the combined data points. Similarly, for the control GP (of the *ℳ*_0_ model) and joint GP (the *ℳ*_1_ model), we have 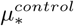 and 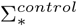 as well as 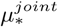 and 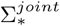, respectively. For the *separate* model, the predictive distribution can be written as the following multivariate normal distribution:

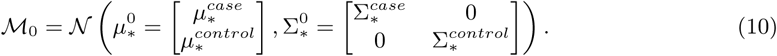

Similarly, the predictive distribution for the *joint* model can also be written as the following multivariate normal distribution:

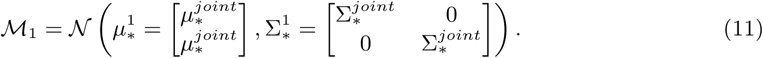

The Kullback-Leibler divergence for any two multivariate normal distributions, say *M*_0_ and *M*_1_, can be computed directly from the formula [Duchi, 2007]

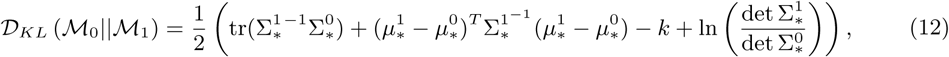

where 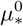 and 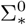 are the parameters of *ℳ*_0_, and 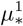 and 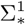 are the parameters of *ℳ*_1_, as discussed above. Also, *k* is the dimension of the multivariate Gaussian, which in our case is 2 *×* 26 (weeks); tr(·) refers to the trace of the matrix; and det refers to the determinant of the matrix.

The symmetric KL divergence gives a KL-score for each probe-set. We specify a threshold for this KL-score such that probe-sets with a KL-score higher than the specified threshold are considered to be differentially expressed. The KL-score can be written as:

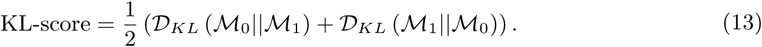

KL-scores do not have a similar interpretation as the Bayes factor. Hence, we empirically set a threshold to identify differential expression prior to an event by taking the mode of the KL-scores of the probe-sets (from all the case-control pairs), which have a BF-score computed from Equation (6) in the range of +*/ −* 1 of the chosen BF-score threshold (in our case, BF-scores in the range 3 to 5 as the threshold is set to 4). The objective of this is to find an appropriate KL-score threshold from the probe-sets that are borderline differentially expressed (or not) according to the BF-scores computed from the whole time-course analysis. Note, however, that a specific value for the threshold is not critical as the pathway level enrichment analysis automatically balances liberal or stringent threshold values.

Lastly, we map the DE probe-sets to their corresponding genes, as explained in Section 3.4. However, in the case of multiple probe-sets mapping to the same gene name, we choose the probe-set with the largest KL-score to represent the gene.

### 3.6 Pathway analysis for personalised differential gene expression results

We propose an empirical hypothesis testing method that can identify statistically enriched pathways from differential probe-set expression analysis results that are computed for all case-control pairs separately as described above in Section 3.4 and Section 3.5 as well as mapped to their corresponding gene names. We define an overall enrichment score for each pathway using the DE genes from each case-control pair using a statistic we call as the *adjusted geometric mean*. Our enrichment analysis uses the number of DE genes from each case-control pair that overlap a given pathway. To account for the fact that a higher number of DE genes in a case-control pair leads to a higher probability of overlap with a pathway, we divide the raw number of DE genes from a case-control pair in a pathway by the total number of DE genes in a case-control pair. Hence, we compute the *scaled pathway overlap f*_*i,j*_ for the *j*^th^ case-control pair and *i*^th^ pathway as

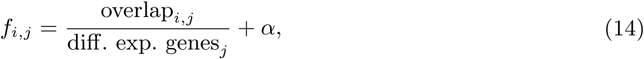

where overlap_*i,j*_ refers to the number of DE genes in the *j*^th^ case-control pair that belongs to the *i*^th^ pathway, diff. exp. genes refers to the number of DE genes in the *j*^th^ case-control pair (assumed to be larger than 0 for all *j*), and *α* is a small constant (*α* = 10^−6^ in our analysis). Assuming *m* case-control pairs, we define an *adjusted geometric mean* for the *i*^th^ pathway as

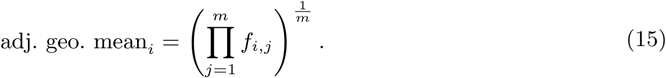

The *adjusted geometric mean* ensures that no case-control pair dominates the overall enrichment score and helps to take into account the different number of DE genes from each case-control pair.

After the *adjusted geometric mean* scores for each pathway are computed, we identify the statistically enriched pathways by performing a permutation test and obtain p-values for each pathway. Let **S** *∈*ℝ^*G×m*^ denote a matrix that stores the BF- or KL-scores from Equations (6) and (13) (where *G* corresponds to the total number of probe-sets and *m* is the number of case-control pairs) such that **S**_*g,j*_ contains the BF- or KL-scores for the *g*^th^ probe-set and the *j*^th^ case-control pair. Our permutation strategy reorders the probe-set labels of the rows, which retains the possible correlations among the scores for the probe-sets across the case-control pairs. In other words, we fix the matrix **S** and shuffle just the associated probe-sets such that each row is randomly assigned a probe-set. After the reordering (shuffling), the probe-sets are again assigned to genes using the same strategy by taking the maximum BF- or KL-scores, and the enrichment scores (*adjusted geometric mean* scores) are computed as above. This process of probe-set label shuffling and computing enrichment scores is repeated 100,000 times to get the permutation distribution that is used to compute the p-values. The permutation distribution acts as the null distribution from which we empirically compute the p-value for a pathway. After this, FDR correction is performed using the Benjamini-Hochberg procedure [Benjamini and Hochberg, 1995] to obtain the set of enriched pathways.

### 3.7 The combined method

We compare our personalised pathway enrichment results with two standard approaches. In the first comparison, we imitate the standard approach of performing DE analysis at the population level and pathway analysis to act as a comparison with our personalised approach. We combine the gene expressions from all the cases and all the controls to obtain a single case-control set of readings, and then compute a list of differentially expressed probe-sets as described above. In this combined method, we again fit two different models. In the first model, we fit two separate GPs, one to case and one to control samples, which we call the *separate* model for the combined method. The second model involves fitting a single GP to the pooled data of all cases and controls, which we call the *joint* model for the combined method. The probe-sets are again mapped to their corresponding genes and in case of multiple probe-sets mapping to a single gene name, we choose the probe-set with the highest BF-score as introduced above. Also, in the rare case that multiple gene names map to a single probe-set, we simply assign the gene name that occurs most often in the annotation database. We perform one-sided Fisher’s exact test to compute p-values [Huang *et al.*, 2009] in order to evaluate the enrichment of each pathway.

In the second comparison, we compare our personalised approach to the results published in [Kallionpää *et al.*, 2014] that correspond to our TC, WSC and WT1D analyses. Briefly, Kallionpää *et al.* [2014] used the rank-product method [Breitling *et al.*, 2004] to identify DE genes. The rank-product algorithm is a rank-statistics based technique for identifying DEGs, where a truly significant gene is expected to appear at the top of independently ranked lists of genes per replicate experiment (e.g. per case) in increasing or decreasing order and score a small geometric mean rank. It is a technique derived from biological reasoning. However, it does not account for the heterogeneity of the disease and it is not suitable for the dynamic analysis of time-course data. For TC analysis, expression values were first normalised for each case-control pair using the z-score and case-wise minimum as well as maximum values are used to identify down- or up-regulated probe-sets. For time-window analyses, in each window (WSC or WT1D), per probe-set fold changes between cases and matched controls were calculated using linear inter-/extrapolation and then used for rank-product analysis. See Kallionpää *et al.* [2014] for further details. In order to keep the pathway level results from [Kallionpää *et al.*, 2014] and our approaches comparable, we performed one-sided Fisher’s exact test (as explained above for combined method) on the gene-level results from all three analyses presented in [Kallionpää *et al.*, 2014] using the pathway information from MSigDB [Subramanian *et al.*, 2005; Liberzon *et al.*, 2015].

## 4 Discussion

The results of this paper demonstrate that a personalised approach of identifying differentially expressed genes (DEGs) and summarising them on a pathway-level can reveal more insight into the progression of heterogeneous diseases, such as type 1 diabetes (T1D), than commonly used non-personalised approaches that assume differences between cases and controls to be consistent across the whole study population, such as the combined method presented in this paper. Even though a significant number of pathways identified by the two approaches overlapped, the combined method was unable to identify the significance of most of the disease-relevant and interesting pathways that were identified by the personalised approach in all the analyses. The combined model identified DEGs in a strict manner that may also be biologically unrealistic, which probably impeded its ability to pinpoint most of the disease-relevant and intriguing pathways.

For validation, the results from the personalised approach were compared to that of the results from Kallionpää *et al.* [2014], who analysed the same datasets using a rank product algorithm introduced by Breitling *et al.* [2004] for identifying DEGs, which cannot account for neither the dynamics of the time-course data nor the heterogeneity. Moreover, they estimated unobserved values in time-window analyses via linear inter-/extrapolation, where we applied Gaussian process modelling, which is known to be more robust. Significant number of pathways identified as enriched by the personalised approach overlapped with theKallionpää *et al.* [2014] results. However, while Kallionpää *et al.* [2014] identified mostly the overarching pathways as enriched, the personalised approach recognised significance of the overarching pathways as well as more specialised pathways that illustrate the intrinsic mechanisms by which the disease develops.

Below, we discuss some of the interesting pathways found enriched by the personalised approach and explore their relevance in terms of T1D as well as the stages of the disease they were found enriched in.

Considering that T1D is a complex autoimmune disease characterised by insulitis, the chronic inflammation of the pancreatic islets of Langerhans caused by autoreactive CD4+ and CD8+ T cells [Clark *et al.*, 2017; Bending *et al.*, 2012; Peakman, 2013; Pugliese, 2017], pathways related to immune response are expected to be enriched along with the T1D pathway. While these particular pathways were not found enriched using the combined model, it did identify interesting and relevant pathways in the TC analysis that largely fall under, but not include, the overarching ‘antigen processing and presentation’ pathway. These were the pathways involving MHC class protein and dendritic cell (DC) maturation. Even though these pathways are highly relevant in the context of the disease, they mostly represent only the initiating events in the development of the disease: release of autoantigens; their uptake by antigen presenting cells (APCs), such as DCs, for antigen presentation in a complex with MHC class proteins [Bending *et al.*, 2012]; and migration of DCs to pancreatic lymph nodes (pLN) to activate *β*-cell specific autoreactive T cells [Bending *et al.*, 2012; Clark *et al.*, 2017], known as DC maturation [Mbongue *et al.*, 2017]. Meanwhile, other important and disease-relevant pathways are underrepresented using the combined model.

The personalised approach also finds the above-mentioned pathways enriched in its analyses, including immune response related and T1D pathways, along with many other disease-relevant pathways. In all the analyses, our approach identifies the pathways related to IFN*γ* signalling and chemokine-mediated signalling as enriched. IFN*γ* is produced by autoreactive CD4+ and CD8+ T cells [Driver *et al.*, 2017] and is believed to play a key role in driving the autoimmune pathogenesis of T1D [Yi *et al.*, 2012; Driver *et al.*, 2017; Souto *et al.*, 2014; Peakman, 2013; Clark *et al.*, 2017; Bending *et al.*, 2012; Borish *et al.*, 2003], even though it is not considered solely a pro-inflammatory cytokine [Driver *et al.*, 2017]. IFN*γ* also results in local up-regulation of chemotactic cues that induce immune cell migration to the islets, for instance via chemokine mediated signalling, where *β*-cells produce certain chemokines that can accelerate or block T1D progression [Clark *et al.*, 2017]. Fascinatingly, our approach also identified a pathway, ‘detection of other organism’, which connotes an existing postulation that environmental factors, such as microbial infections, can trigger the disease process leading to T1D in genetically susceptible individuals [Clark *et al.*, 2017; Bending *et al.*, 2012; Knip and Simell, 2012].

One of the most interesting questions that are asked in T1D studies is regarding the changes that transpire in the time-window leading up to life-changing events, such as seroconversion and clinical onset of T1D. Using the personalised approach, multiple immunologically relevant pathways were revealed to be uniquely enriched in both the time-windows of interest, such as TNF signalling, where TNF-*α* has been linked to the development of T1D [Lee *et al.*, 2005; Bending *et al.*, 2012; Clark *et al.*, 2017; Souto *et al.*, 2014; Borish *et al.*, 2003]; DC differentiation and maturation [Mbongue *et al.*, 2017; Souto *et al.*, 2014]; and cytokine-mediated signalling [Bending *et al.*, 2012; Clark *et al.*, 2017; Peakman, 2013], which acts like an all-encompassing, but vague, pathway for all cytokines. The method was able to determine additional relevant pathways in these two time-windows that were not identifiable by Kallionpää *et al.* [2014] results: immunoglobulin production as well as IL-2 and IL-10 regulating pathways. In fact, it is the increase in production of islet autoantibodies or immunoglobulin that marks the seroconversion event in the life of an individual susceptible to T1D [Kallionpää *et al.*, 2014]. Meanwhile, enrichment of IL-2 and IL-10 signalling pathways before seroconversion indicates the possible anti-inflammatory processes that occur to resist the progression of the disease. IL-10 is an anti-inflammatory cytokine secreted primarily by Tregs and *β*-cell autoantigen recognising CD4+ T cells [Peakman, 2013]. It inhibits the production of multiple pro-inflammatory cytokines, including IFN*γ*, TNF-*α*, IL-5, IL-1*β*, etc. [Borish *et al.*, 2003], and is only marginally less prevalent in T1D patients studied at the time of diagnosis than in healthy subjects [Peakman, 2013]. IL-2 is a cytokine that can lead to prevention or pathogenesis of the disease depending on its own concentration, the concentrations of other local cytokines [Hulme *et al.*, 2012; Hartemann and Bourron, 2012; Pé rol *et al.*, 2016] and polymorphisms in the genes of its pathway [Peakman, 2013]. In low dose, IL-2 signalling is believed to rescue insulin secretion [Hartemann and Bourron, 2012; Pérol *et al.*, 2016]. However, it may result in accelerated autoimmune tissue destruction in the time-window before diagnosis due to the enriched regulation of IL-1 signalling in that time-window as it enhances IL-2 production [Borish *et al.*, 2003; Hartemann and Bourron, 2012].

Our results identify increased number of pathways enriched in the window before T1D onset as compared to the window before seroconversion, demonstrating the mayhem that precedes a clinical diagnosis. Especially, the number of cytokine regulating pathways were increased manifold, where more than half were unique to this time-window. Along with anti-inflammatory cytokines, such as IL-10 and IL-4 [Qiao *et al.*, 2016; Borish *et al.*, 2003; Souto *et al.*, 2014], many pro-inflammatory cytokine regulating pathways were enriched, such as IL-1, IL-1*β*, IL-5 [Borish *et al.*, 2003], IL-6 [Souto *et al.*, 2014; Borish *et al.*, 2003], IL-12 [Mbongue *et al.*, 2017], IL-21 [Bending *et al.*, 2012; Clark *et al.*, 2017; Li *et al.*, 2014], IL-22 [Borish *et al.*, 2003], IFN*γ*, TNF-*α*. In the absence of IFN*γ* and TNF-*α*, cytokines IL-2, IL-1*β* and IL-6 are considered anti-inflammatory [Souto *et al.*, 2014; Clark *et al.*, 2017; Bending *et al.*, 2012; Hartemann and Bourron, 2012; Pérol *et al.*, 2016; Borish *et al.*, 2003], but in their presence, these cytokines aggravate the inflammatory disease pattern, which is probably the case in the time time-window before T1D diagnosis.

Some of the pathways that were found enriched in the time-window before T1D diagnosis were also found enriched during the early stages of T1D progression using the personalised approach, possibly indicating that key players from late stages of the disease may already be detected at the early stages. These included both pro- and anti-inflammatory pathways, such as those of IL-1 and IL-4 as well as IFN*α* and PD-1 signalling. IL-1 is a pro-inflammatory cytokine that enhances the production of IL-2, encourages B cell proliferation, and increases immunoglobulin production [Borish *et al.*, 2003; Hartemann and Bourron, 2012]; whereas IL-4 is an anti-inflammatory Th2 cytokine that inhibits autoimmunity by down-regulating the production of pro-inflammatory cytokines, such as IL-1, IL-6 and TNF-*α* [Borish *et al.*, 2003; Souto *et al.*, 2014; Qiao *et al.*, 2016]. Through mice studies, IFN*α* and PD-1 signalling pathways have been established as important contributors to T1D pathogenesis from an early stage of the disease [Marro *et al.*, 2017; Li *et al.*, 2008; Mbongue *et al.*, 2017; Martinov *et al.*, 2016]. Where up-regulation of IFN*α* in pLN is an initiator of the pathogenesis [Li *et al.*, 2008], up-regulation of programmed cell death protein 1 (PD-1) signalling prevents T1D and promotes self-tolerance by suppressing the expansion and infiltration of autoreactive T cells in the pancreas [Granados *et al.*, 2017; Martinov *et al.*, 2016; Mbongue *et al.*, 2017]. In fact, blocking IFN*α* signalling before clinical T1D onset has been shown to prevent *β*-cell apoptosis or even abort T1D progression [Marro *et al.*, 2017]. Additionally, PD-1 pathway has been proposed as a target for novel therapy for preventing and modulating autoimmunity [Granados *et al.*, 2017].

Fascinatingly, natural killer (NK) cell mediated cytotoxicity pathway was found to be uniquely enriched during the early stages of T1D. NK cells are believed to be involved in multiple steps of the immune-mediated attack causing T1D as they are known to interact with antigen-presenting T cells, secrete pro-inflammatory cytokines and induce apoptosis in the target cells [Rodacki *et al.*, 2006; Qin *et al.*, 2011]. Similarly, Fas signalling pathway was found to be uniquely enriched before seroconversion. Since it is one of the pathways mediated by autoreactive CD8+ T cells that is directly involved in the destruction of *β*-cells [Bending *et al.*, 2012; Clark *et al.*, 2017; Mbongue *et al.*, 2017], it demonstrates that *β*-cell killing can be observed much before the clinical onset of T1D.

Even though the personalised approach is able to identify many immunologically- and disease-relevant pathways, it has scope for further development. The current implementation assumes Gaussian distributed data; it may be possible to improve the accuracy of differential gene expression detection for data sets that have notably non-Gaussian characteristics either by using a different likelihood model or by performing appropriate transformations. In addition, the proposed approach has been implemented for a matched case-control setting. However, with small modifications to the model, it could be extended to a non-matched case-control setting, where each case is compared to all the controls in the dataset.

## 5 Acknowledgements

This work has been supported by the Academy of Finland (grants no. 292660) and Business Finland. We would like to acknowledge the computational resources provided by Aalto Science-IT and CSC-IT Center for Science, Finland. We would also like to thank Markus Heinonen for his valuable insights.

## 6 Author contributions

J.S., S.R. and H.L. co-developed the method presented in this paper. S.R. implemented the method and J.S. interpreted the results as well as supervised S.R. H.L. oversaw the whole project and supervised J.S. and S.R. All authors contributed to the writing of this manuscript.

