## Supplementary Figures for "A personalised approach for identifying disease-relevant pathways in heterogeneous diseases"

Juhi Somani*, Siddharth Ramchandran* and Harri Lähdesmäki

Department of Computer Science,
Aalto University,
02150 Espoo, Finland.
``

### 1 Supplementary figures

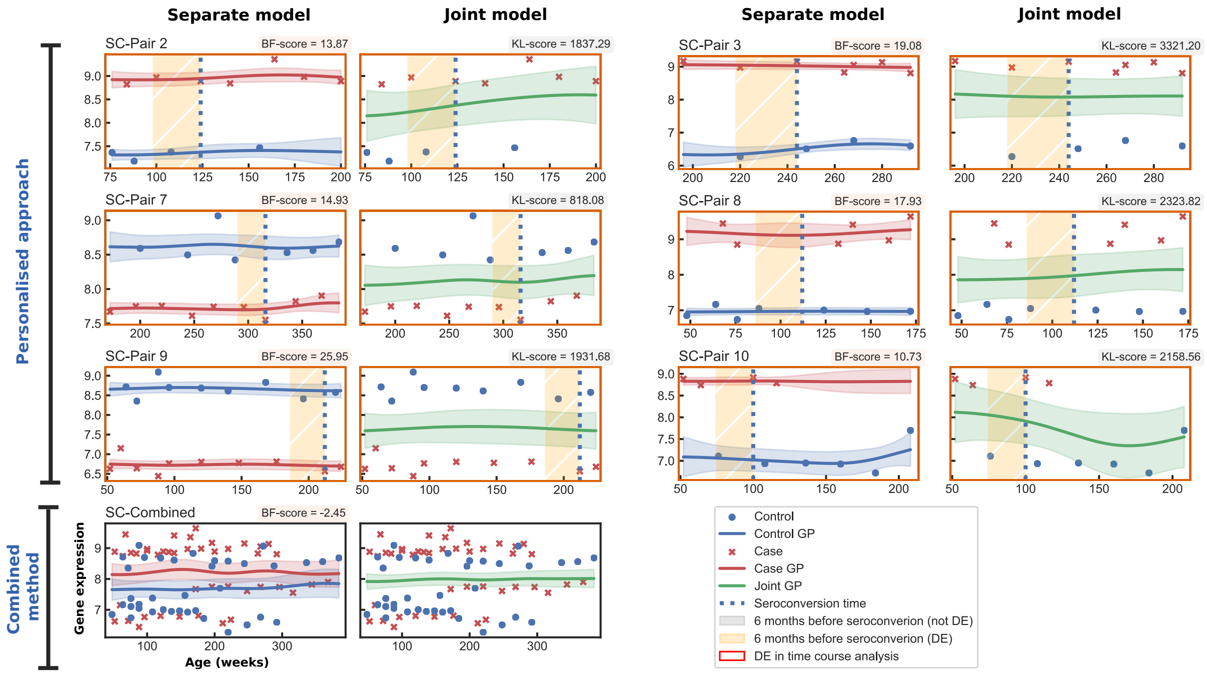


Figure 1: Gene expression plots for the *IRF5* gene, visualising the GP model fittings of the separate and joint models for the six case-control pairs from **Dataset 1**. A red border around a plot signifies differential expression (DE) in the time-course analysis and an orange shaded window signifies DE in the time-window analysis. Here, pairs from **Dataset 1** are prefixed with 'SC-'. All profiles belong to the same probe-set as all pairs, including the combined method, identified the same probe-set to have the largest BF-score for *IRF5*.

*Co-first author; contributed equally to this work

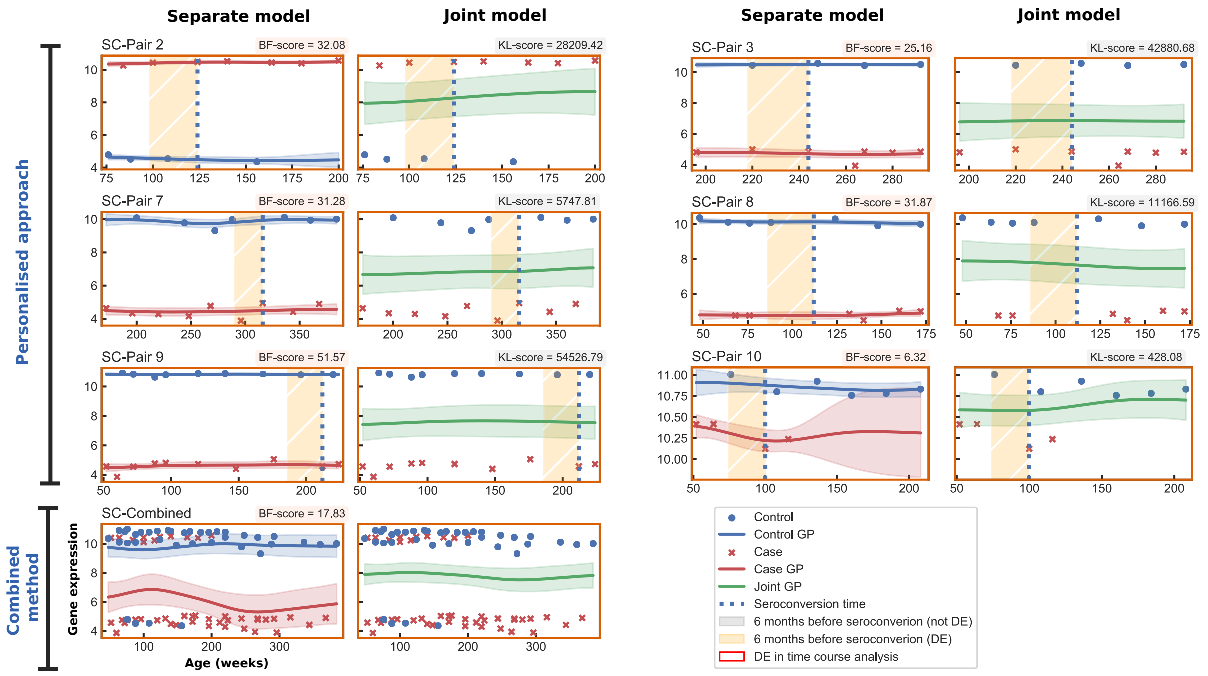


Figure 2: Gene expression plots for the *HLA-DPB1* gene, visualising the GP model fittings of the separate model and joint model for the six case-control pairs from **Dataset 1**. A red border around a plot signifies differential expression (DE) in the time-course analysis and an orange shaded window signifies DE in the window analysis. Here, pairs from **Dataset 1** are prefixed with 'SC-'. All the visualisations belong to the same probe-set, including in the combined method.

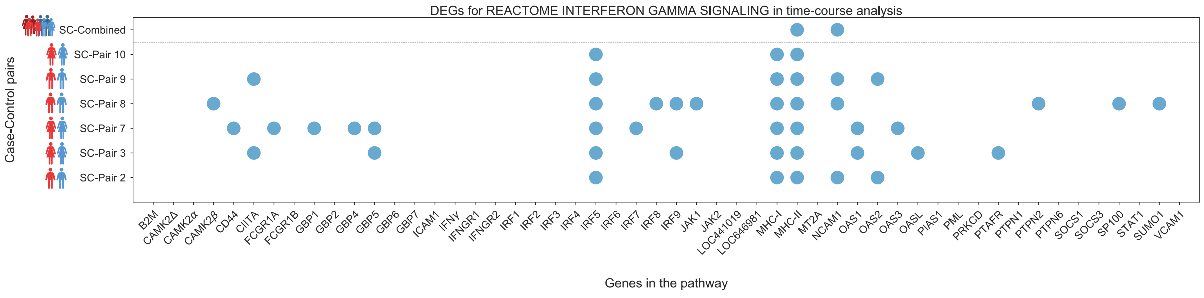


(a)

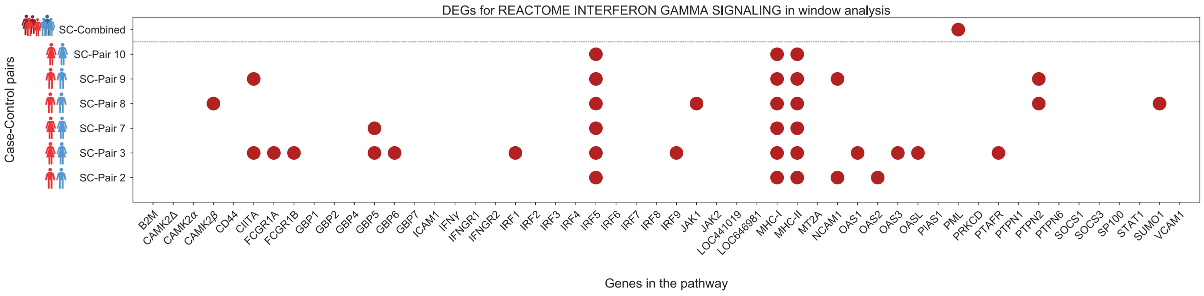


(b)

Figure 3: A comparative visualisation of the DEGs between the two approaches for the reactome interferon gamma signalling pathway using **Dataset 1**, prefixed with 'SC-': (a) TC analysis and (b) WSC analysis. A coloured dot signifies that the gene is DE in the corresponding case-control pair. MHC classes I and II are independently marked as DE for each case-control pair if at least one probe-set in the respective group of HLA genes is detected as differentially expressed.

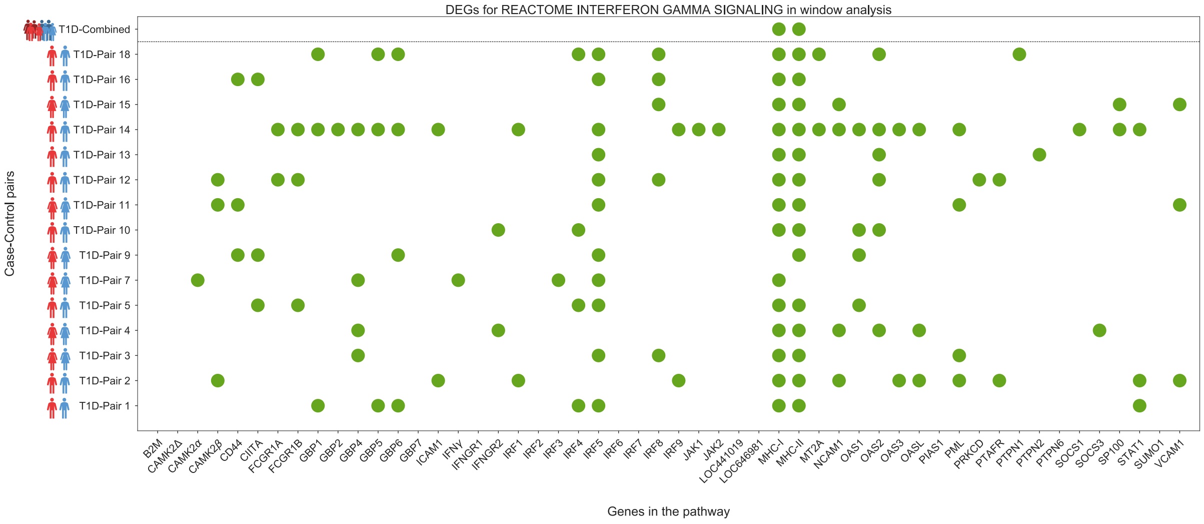


Figure 4: A comparative visualisation of the DEGs between the two approaches for the reactome interferon gamma signalling pathway pathway using **Dataset 2**, prefixed with 'T1D-', in the WT1D analysis. A coloured dot signifies that the gene is DE in the corresponding case-control pair. MHC classes I and II are independently marked as differentially expressed for each case-control pair if at least one gene in the respective group of HLA genes is detected as differentially expressed.
